# Viable But Non-Culturable Cells Are Persister Cells

**DOI:** 10.1101/186858

**Authors:** Jun-Seob Kim, Nityananda Chowdhury, Thomas K. Wood

## Abstract

Bacteria have two dormant phenotypes: the viable but non-culturable (VBNC) state and the persister state. Both resting stages arise without mutation and both have been linked to chronic infections; however, persister cells revive rapidly whereas the cell population called VBNC is reported to not resuscitate. Here we investigated the relatedness of the two stress-induced phenotypes at the single-cell level by using transmission electron microscopy and fluorescent microscopy to examine cell morphology and by quantifying cell resuscitation. Using the classic starvation conditions to create VBNC cells, we found that the majority of the remaining *Escherichia coli* population are spherical, have empty cytosol, and fail to resuscitate; however, some of the spherical cells under these classic VBNC-inducing conditions resuscitate immediately (most probably those with dense cytosol). Critically, all the culturable cells became persister cells within 14 days of starvation. We found that the persister cells initially are rodlike, have clear but limited membrane damage, can resuscitate immediately, and gradually become spherical by aging. After 24 h, only rod-shaped persister cells survive, and all the spherical cells lyse. Both cell populations formed under the VBNC-inducing conditions and the persister cells are metabolically inactive. Therefore, the bacterial population consists of dead cells and persister cells in the VBNC-inducing conditions; i.e., the non-lysed particles that do not resuscitate are dead, and the dormant cells that resuscitate are persister cells. Hence, “VBNC” and “persister” describe the same dormant phenotype.

## INTRODUCTION

Bacteria have elegant ways to survive during stress^1,2^, such as that arising from inevitable nutrient depletion as well as antibiotic exposure, and two distinct phenotypes have been described in which the cells enter a non-heritable, reversible, dormant state: viable but non-culturable (VBNC) cells^3^ and persister cells^4^. In what have become known as persister cells, Hobby et al.^4^ determined in 1942 that 1% of *Staphylococcus aureus* cells are not killed by penicillin and that these antibiotic-tolerant cells are metabolically inactive. Forty years later, VBNC cells were first described as those *Escherichia coli* and *Vibrio cholerae* cells that are present after an extended period (two weeks) in salt water microcosms that are not culturable on selective and non-selective media upon which they are usually capable of growth^3^; however, a few stimuli such as nutrients and temperature shifts serve to resuscitate VBNC cells^5^.

The two states of dormancy have much in common. Both persisters^6^ and VBNCs^5^ have been linked to chronic infections, both occur in biofilms^5,7^, and both cell types have been generated by more than one kind of stress; for example, oxidative and acid stress^5,8^. Furthermore, the genetic basis for both cell types is not well characterized. For persisters, toxin/antitoxin systems^9,10^, the alarmone guanosine tetraphosphate (ppGpp)^11^, and the stationary-phase sigma factor^8^ RpoS clearly play a major role in their formation; however, many systematic studies such as transposon-sequencing^12^, protein expression^13^, and gene knockouts^14,15^ have not yielded significant additional insights into persister cell formation, and it appears any toxic protein that slows bacterial growth induces persistence, even in the absence of ppGpp^16^. For VBNC cells, RpoS^17,18^, the transcriptional regulator OxyR that controls genes related to oxidative stress^5^, and toxin/antitoxin systems^19^ have been linked to VBNC cells. Hence, it has been suggested that these two survival states may be part of a “dormancy continuum”^19^; i.e., the two kinds of resting states may be related with VBNCs as the more dormant of the two states. The key feature that distinguishes persister cells from VBNC cells is that VBNC cells cannot be resuscitated under normal conditions while persister cells can be easily converted to normal cells that are sensitive to antibiotics or other stresses^19^.

VBNC cells and persister cells share many similar features, and they co-exist^20-23^; however, no studies have been performed to compare these two stages of dormancy based on their physiology and morphology. Here, we demonstrate that viable fraction of VBNC cells generated from nutrient depletion are persister cells by comparing their antibiotic tolerance, rates of resuscitation, morphology, and metabolic activity. The remainder of the VBNC cell fraction are dead. Hence, the dormant cell phenotype known as VBNC is the same as that known as persister cells.

## RESULTS

We hypothesized that VBNC cells are related to persister cells, so we generated VBNC cells using long-term nutrient depletion (prolonged exposure to 0.85% NaCl buffer), a well-known method to form VBNC cells^24,25^, and studied the cell population under these VBNC-inducing conditions in a temporal fashion (over 7 weeks) by measuring the number of viable, culturable and persister cells as well as by characterizing their morphology. Total cells and viable cells were determined using a hemocytometer with Syto 9 and PI staining, which stains live and dead cells, culturable cells were determined by colony forming units (CFU) without antibiotic treatment, and persister cells were determined by counting CFU after antibiotic treatment. To enable a comparison to the cell population in the VBNC-inducing conditions, we generated persister cells in high numbers by pre-treating cells with rifampicin^26^. We have verified seven ways that these rifampicin pre-treated cells are persister cells (muti-drug tolerance, easy conversion from persister state to non-persister state, metabolic dormancy, no change in the minimum inhibitory antibiotic concentration compared to exponential cells, no resistance phenotype, similar morphology to natural persisters, and similar resuscitation as natural persisters) (manuscript in revision). We also compared the rates at which individual cells resuscitate by using agarose pads and light microscopy as well as compared cell morphology via TEM.

### The small, culturable fraction formed under VBNC-inducing conditions are persister cells

In an *E. coli* population with an initial size of 2 x 10^8^ total cells/mL (viable + non-viable cells) (Fig. 1) as determined using a hemocytometer to ensure all particles were counted (Supplementary Fig. 1), we found that the number of culturable cells decreased dramatically from 10^8^ to 10^3^ cells/mL in seven weeks (Fig. 1). Critically, the number of persister cells (i.e., those that survive antibiotic treatment) increased from 10^3^ to 10^5^ cells/mL in two weeks (Fig. 1), and all the culturable cells under the VBNC-inducing conditions become persister cells; hence, all the culturable cells under the VBNC-inducing conditions became persister cells and remained persister cells for seven weeks (when the experiment was stopped). This result indicates that starvation induces persister cell formation and that all the culturable cells in the VBNC-induced condition phenotypically behave exactly like persister cells after two weeks.

**Fig. 1.**
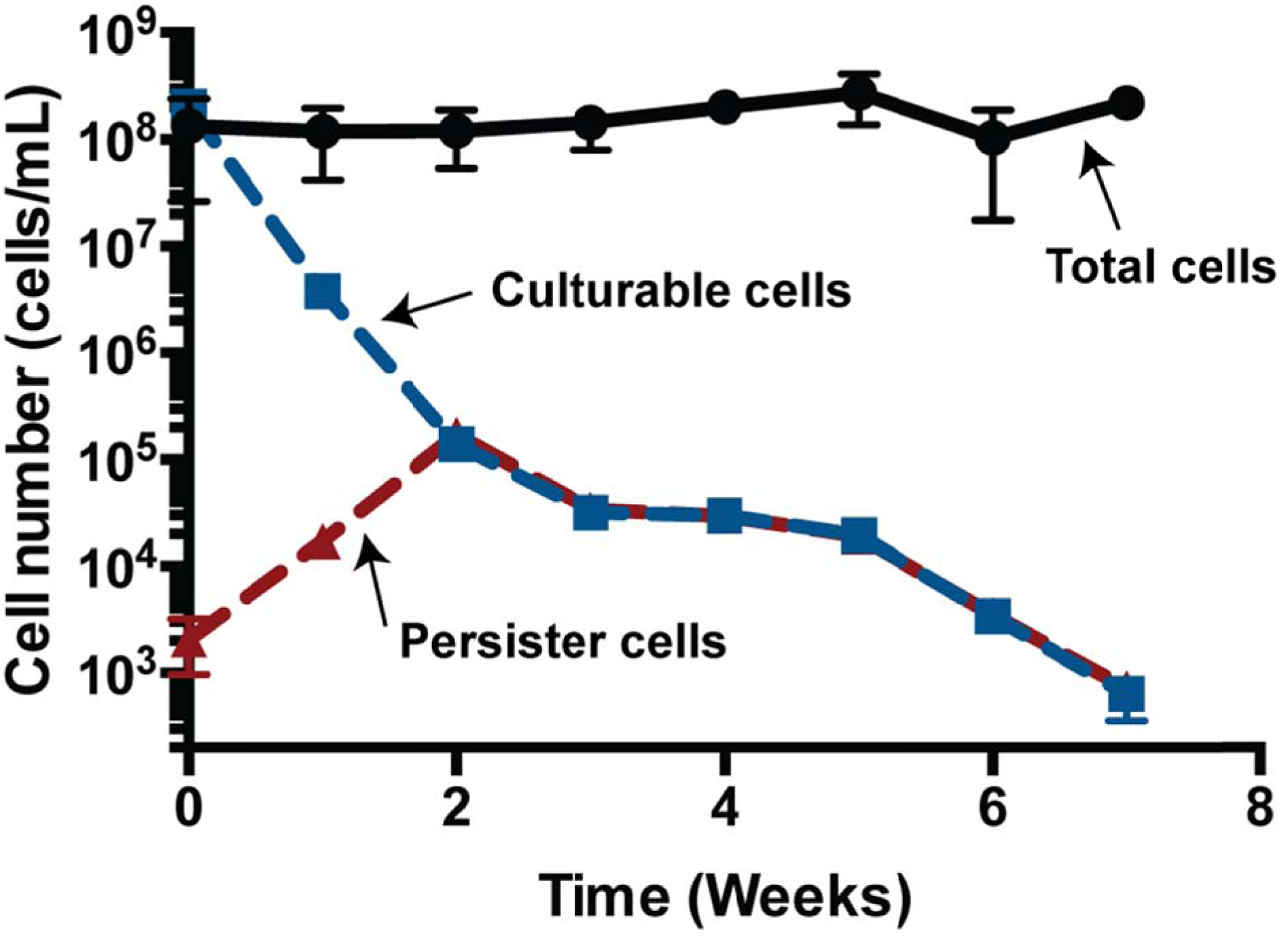
Temporal change in VBNC culturable and persister cells. The total live cell population is indicated by black circles and was determined via by hemocytometer with Syto 9 and PI staining. Total cells indicates cell numbers as visualized by membrane stain Syto 9. The cell population stained by PI was less than 2% of the total cell number. The culturable cell number is indicated by blue squares and was determined by colonies formed on plates. Persister cells are indicated by red triangles and were determined by colonies formed on plates after ampicillin treatment for 3 h. VBNC cells were treated with ampicillin for 3 h.

### The small, culturable fraction formed under VBNC-inducing conditions resuscitate like persister cells

Persister and VBNC cells share many features like toleration of many stresses (e.g., antibiotics, heat, acid)^19^, induction by various environmental stresses^19,27^, and dormancy in stressful environments^3,28^. However, unlike what has been reported for VBNC cells, persister cells resuscitate easily by nutrients when the stress is removed^19^. To confirm that the live cell population created under the VBNC conditions, that we found to be antibiotic tolerant (Fig. 1), resuscitate like persister cells, the resuscitation rate of individual cells on agarose pads was determined.

The positive control, exponential cells, began cell division immediately (Supplementary Video 1), which indicates, that there are no delays or artifacts affecting growth inherent with our agarose pad method. The persister cells resuscitated with various waking times (0 to 6 h) (Supplementary Video 2), and had a similar growth rate as exponential cells, which means that persister cells convert to normal growing cells upon waking (**data not shown**). In contrast to persister and exponential cells, on the agarose pads, most of the cells under the VBNC-inducing condition were small, not dense, and spherical (Supplementary Video 3). After 5 weeks, the total number of viable cells in the population grown under the VBNC-inducing conditions did not change appreciably compared to 0 to 4 weeks, which indicates that this morphologically abnormal cell population is the major cell phenotype. Critically, this cell spherical phenotype found under the VBNC-inducing conditions did not resuscitate in 72 h on the agar pads (Supplementary Video 3).

Since the number of persister cells formed under the VBNC-inducing conditions was very low after 5 weeks (about 10^4^ cells/mL, Fig. 1), 2 week old VBNC cultures with ampicillin pre-treatment were examined, and it was found that some spherical cells revived immediately by becoming rod-shaped then dividing (Supplementary Fig. 2, blue arrows) while some did not revive (Supplementary Fig. 2, red arrows). Similarly, rods formed under the VBNC-induced condition revived immediately (Supplementary Fig. 2, green arrows).

### The small, culturable fraction formed under VBNC-inducing conditions and persister cells have the similar morphology

Cells formed under VBNC-inducing conditions cells are frequently spherical^29,30^, exponential-phase *E. coli* cells are rods; and persister cells have not been characterized well via microscopy due to the difficulty in getting populations with significant fractions of persister cells. To explore further the relationship between the cell population formed under VBNC-inducing conditions and persister cells, we utilized TEM. The TEM images confirm that exponentially-growing cells (positive control) are rod shaped and healthy (i.e., dense cytosol, Fig. 2); hence, healthy cells have dense cytosol with this visualization method^31^. Critically, the majority of the cells (5 weeks) formed under VBNC-inducing conditions have empty cytosol, and they have either intact or damaged membranes (Fig. 2, Supplementary Fig. 3). The empty region in the cytosol might explain why most VBNC cells do not resume normal growth: they are dead, since empty cytosol is a sign of cell death^32,33^. Using the TEM images, the viability of cells formed under the VBNC-inducing condition was estimated by counting cells with dense cytosol as live cells and cells with empty cytosol as dead: from our images, 99.5% of the total VBNC cell-like particles have the characteristics of dead cells (Table 1).

**Fig. 2.**
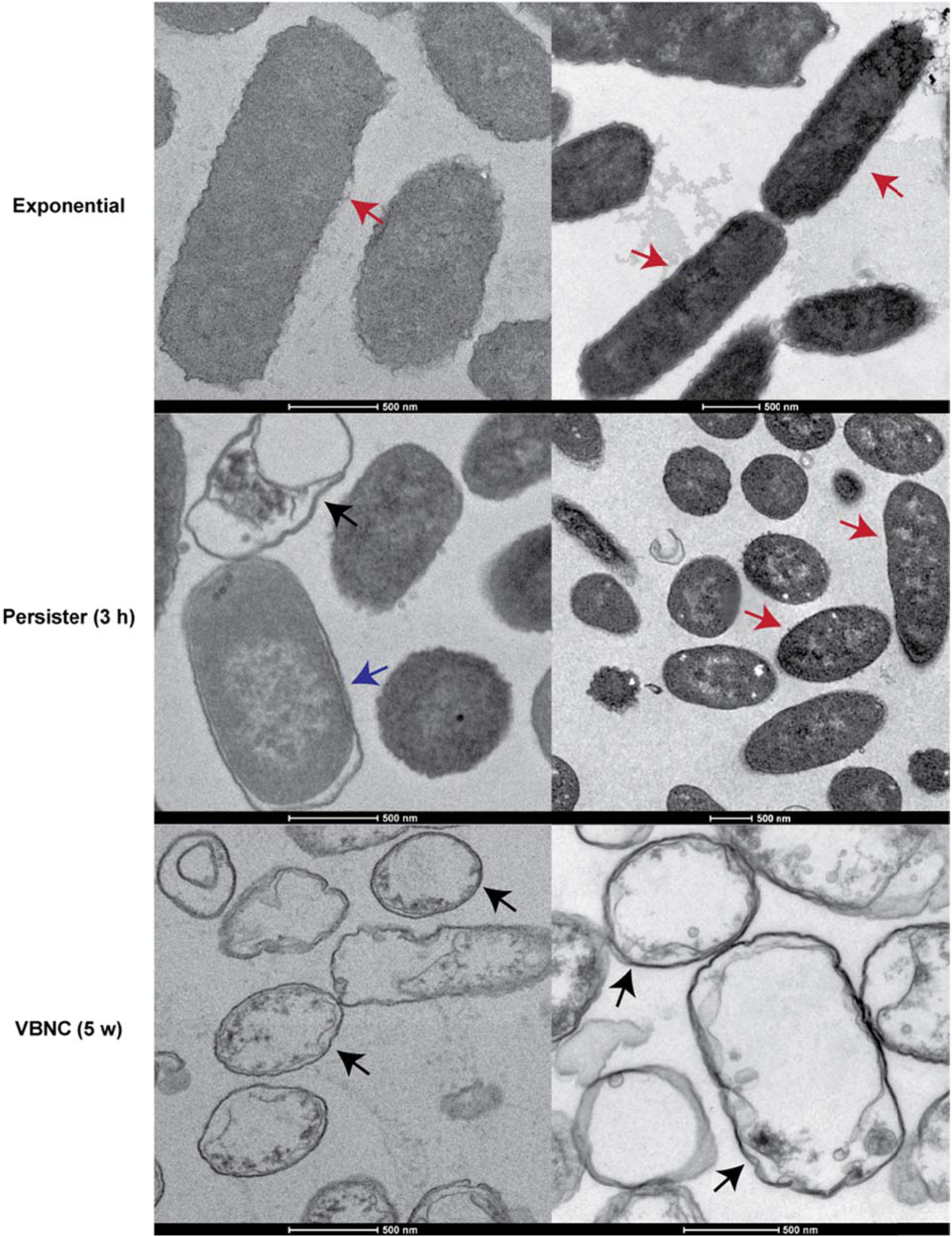
TEM images of exponential, persister, and VBNC cells. Exponentially-grown cells, rifampicin-induced persister with 3 h ampicillin treatment, and 5 week (5 w)-old VBNC cells (ampicillin-treated for 3 h) in saline solution were visualized by TEM. Red arrows indicate cells with dense cytosol and intact membranes, blue arrows indicate cells in the intermediate stage of material loss, and black arrows indicate cells with empty cytosol but intact membranes. Scale bar indicates 500 nm.

**Table 1.**
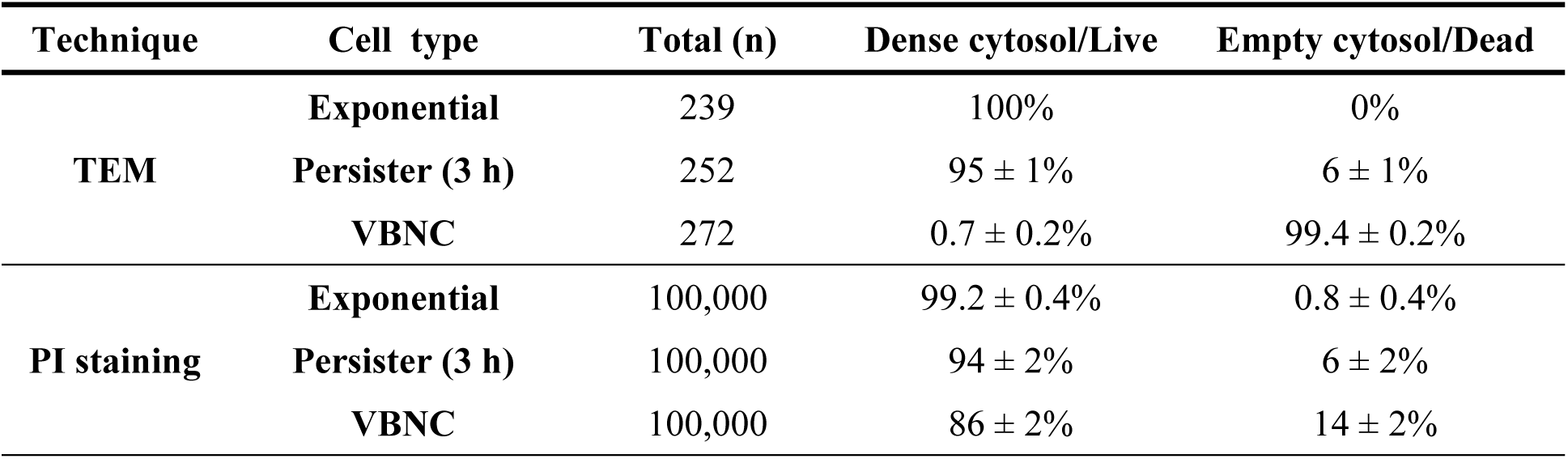
Comparison of the viability of each cell types via TEM and PI staining method.

| **Technique** | **Cell type** | **Total (n)** | **Dense cytosol/Live** | **Empty cytosol/Dead** |
| --- | --- | --- | --- | --- |
| **TEM** | **Exponential** | 239 | 100% | 0% |
|  | **Persister (3 h)** | 252 | 95 ± 1% | 6 ± 1% |
|  | **VBNC** | 272 | 0.7 ± 0.2% | 99.4 ± 0.2% |
| **PI staining** | **Exponential** | 100,000 | 99.2 ± 0.4% | 0.8 ± 0.4% |
|  | **Persister (3 h)** | 100,000 | 94 ± 2% | 6 ± 2% |
|  | **VBNC** | 100,000 | 86 ± 2% | 14 ± 2% |

TEM images of younger cells (2 weeks) formed under the VBNC-inducing conditions showed two types of spherical cells: those with dense cytosol and those with empty cytosol (Supplementary Fig. 4). This suggests that the spherical cells that do not resuscitate are dead (empty cytosol), whereas spherical cells with dense cytosol resuscitate.

Newly-formed persister cells (3 h) are also rod shaped, but a few of them are clearly injured with damaged membranes (Fig. 2, Supplementary Fig. 5). Additionally, some of the persister cells are spherical after 3 h. Critically, a few of the VBNC-like (spherical/empty cytosol) shaped cells were also present in the persister cell population after 3 h (Fig. 2) This result suggests that perhaps the newly-formed persister cells undergo a morphological change as they age; hence, we monitored the shape of persister cells via fluorescent microscopy with FM 4-64 staining (which helped visualize the cells). As expected, the exponential cells were cylindrical (positive control) but rifampicin-induced persister cells had shorter cell lengths after 3 h (Fig. 3A); this reduction in cell volume increased with age for persister cells (Fig. 3A). Based on this observation, cell roundness was calculated. The average roundness, which indicates how closely the shape of a cell approaches that of a circle, of exponential cells was 0.4 while the average roundness of persister cells was 0.6 after 3 h (Fig. 3B); hence, persister cells were more spherical than exponential cells and began to appear more similar to VBNC cells as they aged. The spherical shape for the persister cells from 3 to 9 h increased as evidenced by the shift to the right (increasing roundness) of the cell population average (Fig. 3B).

**Fig. 3.**
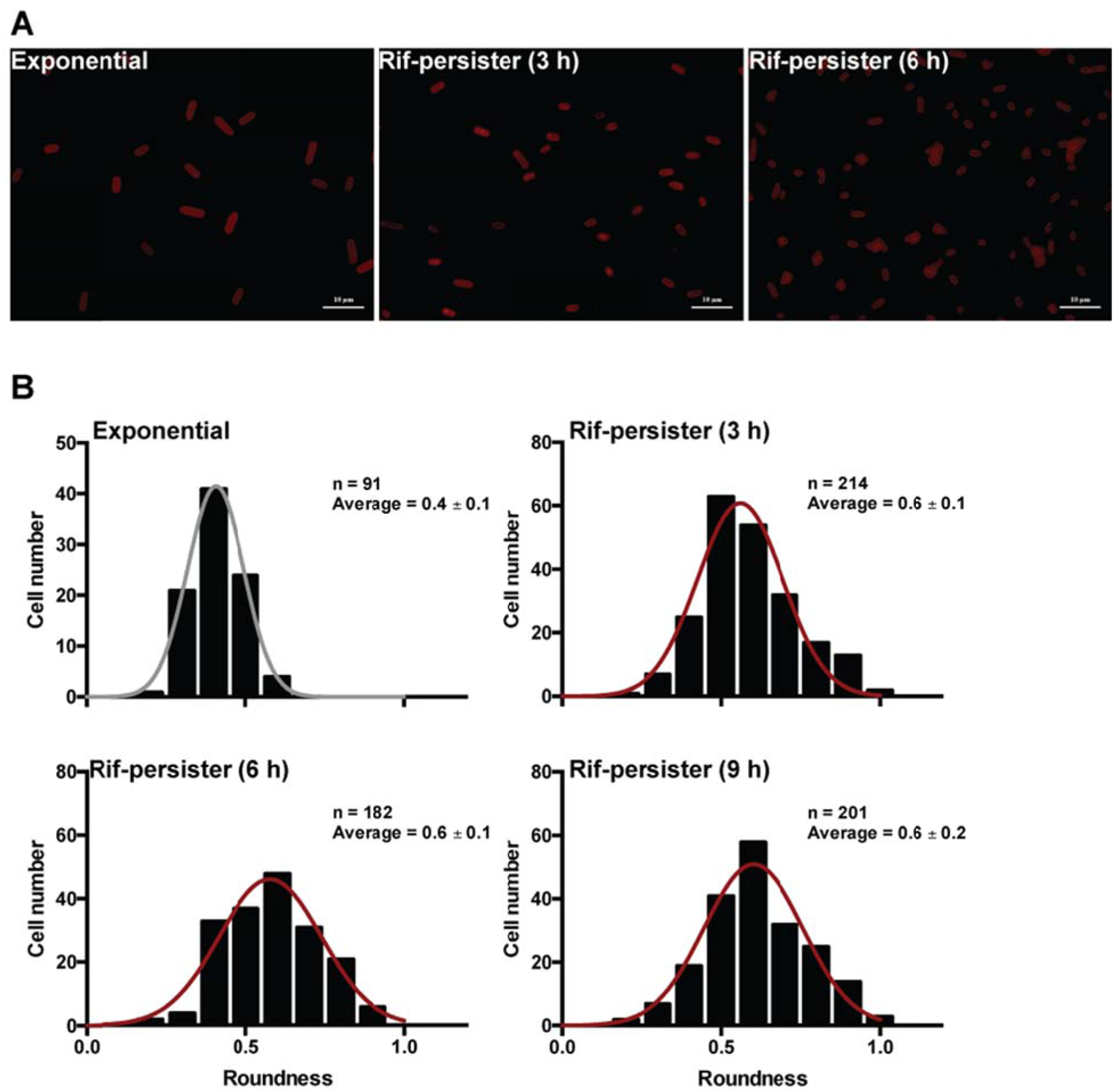
Aging persister cells become spherical. Rifampicin-induced persister cells in LB in the presence of 100 μg/mL of ampicillin were observed after 3, 6, 9, and 12 h via fluorescence microscopy with FM 4-64 staining. (**A**) Representative microscopic images for exponentially-growing cells and aging persister cells. (**B**) Calculation of cell roundness by ImageJ. Compared to exponential cells, persister cells have increasing roundness with age (distribution of roundness shifts to right.

After 24 h, rod-shape persister cells remained but there was the accumulation of cell membrane debris as visualized by FM 4-64 staining since the population of persisters usually decreases (Supplementary Fig. 6); hence, the persister cells that became spherical died and lysed, much like the cells formed under the VBNC-inducing conditions as demonstrated by TEM (Fig. 2). We hypothesized that the empty spheres in the persister culture were dead cells and only the rod-shape persister cells could resuscitate; i.e., cells in the persister cell culture slowly die over 24 h in the presence of ampicillin. We tested this hypothesis by measuring the viability of persister cells and found that in 24 h, the viable persister cell fraction decreased by 70-fold (Supplementary Fig. 7), which is much slower than the death rate of exponential cell by ampicillin (10,000-fold lower than exponential cell death in 24 h). Therefore, cells in the persister population die over 24 h.

TEM images of fresh persister cells (3 h) (Supplementary Fig. 5) and old persister (24 h) cells (Supplementary Fig. 8) showed both the 3-h and 24-h persister cells were smaller compared to exponentially-growing cells (Fig. 2) which agrees with FM 4-64 results (Fig. 3). Moreover, the spheres seen in the fresh persister population had dense (full) cytosol. Since, most of the old persister cells (24 h) are spherical with empty cytosol (Supplementary Fig. 8), they resemble cells of the VBNC cultures (Supplementary Fig. 3 and Supplementary Fig. 4); however, some rod-shaped cells with full cytosol remain in the 24 h persister culture (Supplementary Fig. 8), and it these rod-like cells that remain after 24 h that were seen to revive on agarose pads like fresh persister resuscitation (Supplementary Fig. 9). Hence, persister cells that are aged in the presence of antibiotic appear the same as cells formed under the VBNC-inducing conditions treated with ampicillin.

### The small, culturable fraction formed under VBNC-inducing conditions and persister cells are metabolically inactive

VBNC cells^34,35^ and persister cells^36,37^ are both dormant. To investigate the metabolic activity of the cells formed under VBNC-inducing conditions and persister cells used in this study, redox sensor green, which indicates cellular metabolic activity, and PI, which indicates membrane damage, were utilized with flow cytometry so that the signals could be quantified. In terms of metabolic activity, both the 5-week-old cells formed under the VBNC-inducing conditions and the 3-h persister cells had lower metabolism than the dead cell control (Fig. 4A), which indicates that they do not have any metabolic activity (perhaps the persister cells and cells formed under VBNC-inducing conditions have lower metabolic signal than the dead cells since their membrane structures are altered as we showed with TEM). As a positive control, there was high metabolic activity with the exponential cells, as expected (Fig. 4A), and unstained exponential cells (negative control) had the expected low signal (Fig. 4A).

**Fig. 4.**
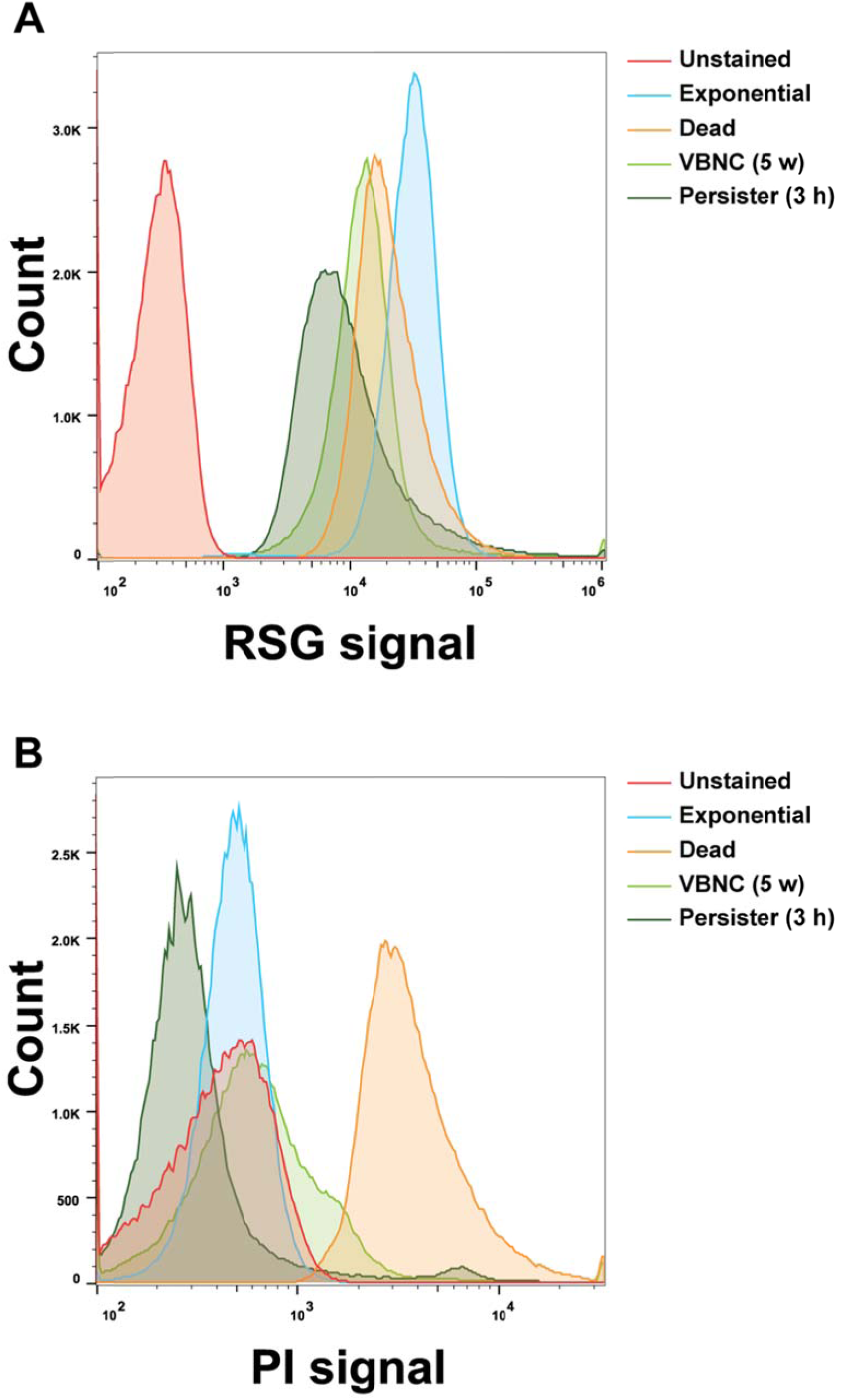
Metabolic activity of persister and VBNC cells. Metabolic activity of exponentially-growing cells, persister cells (3 h), and VBNC cells (5 weeks: “5 w”, ampicillin-treated for 3 h) as determined by the BacLight RedoxSensor Green (RSG) Vitality Kit and flow cytometry. (**A**) Metabolic activity of each cell culture based on the RSG signal. (**B**) Cell viability of each cultures based on PI signal. As controls, unstained exponential cells and dead cells (70% isopropanol treatment for 1 h) were employed.

As with the fluorescence microscopy data where PI staining indicated little change in the total number of live cells (Fig. 1), only 15% of cell population in the VBNC culture appeared dead using PI staining (Fig. 4B). However, the TEM results indicate that a majority cells (>99%) formed under the VBNC-inducing conditions have lost their cellular mass and are dead (Supplementary Fig. 3). As controls, the dead cell fraction of exponentially-growing cells was 0.5% and the fraction of dead cells in the isopropanol-treated dead cell control was 99.3% (Fig. 4B). These conflicting results of the TEM images showing nearly all the cells are dead (i.e., have empty cytosol) and PI staining indicating most cells are viable indicate that PI staining works in many situations but unfortunately, not for VBNC cultures where it has been used previously^38,39^.

## DISCUSSION

Although many similarities exist in the published descriptions of VBNC and persister cells (the main distinction is that VBNC cells are not culturable), these two phenotypes have not been compared experimentally. By assaying for persister cells in the VBNC population, we discovered here that the viable cells formed under the VBNC-inducing conditions are persister cells (tolerant to antibiotics) (Fig. 1); hence, persister cells may be formed from nutritive stress, and the culturable fraction of the cells formed under the VBNC-inducing conditions are persister cells. For the cell population previously known as VBNC, most of the cells are dead as evidenced by their empty cytosol (Fig. 2 and Supplementary Fig. 3). Additionally, we also found that there are some spherical cells in the persister culture after 3 h that lack cellular material (Fig. 2 and Supplementary Fig. 5), and that persister cells become spherical as they age and have an increasing fraction of cells with empty cytosol (Fig. 3 and Supplementary Fig. 8). Furthermore, persister cells die over 24 h (Supplementary Fig. 5, 6, & 7). Hence, cultures of 24 h-old persisters and cells formed under VBNC-inducing conditions have the same morphology and both cultures have live cells (dense cytosol) that are found in a background of dead cells, and it is theses intact *E. coli* cells that are able to resuscitate. We also found both the culturable fraction of the cells formed under VBNC-inducing conditions and persister cells have no metabolic activity (Fig. 4) and can resuscitate immediately. Therefore, the viable cells formed under the VBNC-inducing conditions appear to be the same as persister cells based on antibiotic tolerance, morphology, resuscitation rates, and metabolic activity. We suggest then that the terms VBNC and persistence describe the same phenotype for dormant cells and that the term VBNC should be replaced with persister cells since VBNC cells do not represent a separate cell phenotype.

One non-intuitive finding here is that most of the “VBNC” cells shrink and are dead since they contain little cytosolic content (i.e., proteins, Fig. 2); yet, they do not stain with PI, which indicates their membranes are not visibly damaged. This lack of staining by PI has led many groups to surmise that these cell remnants are alive and are difficult culture; however, TEM clearly indicates they are dead. What remains unclear is how the cellular contents are lost without damaging the membranes. The loss of cell content does fit well with the fact that others have found that cells shrink and become spherical (e.g., *Salmonella typhi*^29^, *Edwardsiella tarda*^30^) as we have demonstrated with aging persister cells (Fig. 2 and 3). Perhaps this loss of cell content occurs as the persister cells age through blebbing as seen in the presence of β-lactams^40^ which would keep the membrane remains intact. Regardless of the aging mechanism, stressed cells die and the remaining viable fraction are persister cells.

## METHODS

### Bacterial strain and growth conditions

*E. coli* K-12 BW25113^41^ was used in this study. All experiments were conducted at 37 °C in saline (0.85% NaCl) for VBNC cultures and lysogeny broth (LB)^42^ for persister cells.

### Total, viable, and antibiotic-tolerant cells in VBNC cultures

To generate VBNC cells, overnight LB cultures were inoculated into 25 mL fresh LB using a 1:1000 dilution and incubated until the turbidity at 600 nm reached 3. Cells (5 mL) were harvested by centrifugation at 3,500 x g, 4 °C for 10 min, and the cell pellet was diluted 1:10 into saline (0.85% NaCl, 50 mL) and incubated at 37 °C, 250 rpm. Every week, cell viability and total cell counts were determined with a hemocytometer after staining with Syto9 (Molecular Probes, Eugene, OR, USA) and propidium iodide (PI, Molecular Probes, Eugene, OR, USA) and visualizing cells with a fluorescence microscope (Zeiss Axioscope.A1) with 400x magnification (40x, 10x), bright field, FITC and PI channel (exposure time ~2,000 ms). The images were superimposed and counted manually. The culturable cell number in the VBNC populations were determined by counting colonies on LB agar plates after 16 h. To determine the number of antibiotic-tolerant cells (i.e., persister cells) in the VBNC population, 25 mL were harvested, the cell pellet was resuspended into 25 mL of LB containing ampicillin (100 μg/mL) for 3 h to remove the antibiotic-sensitive cells (which lyse)^10^, centrifuged, washed by phosphate buffered saline (PBS) twice, and plated onto LB agar for 16 h.

### Persister and VBNC cell resuscitation

To generate persister cells, 25 mL of exponentially-growing cells (turbidity at 600 nm ~0.8) were treated with rifampicin (100 μg/mL) for 30 min^26^ and harvested by centrifugation (3,500 x g, 4 °C for 10 min). The cell pellets were resuspended in 25 mL of LB containing ampicillin (100 μg/mL) and incubated for 3 h to remove non-persisters via lysis.

For resuscitation, 1 mL was taken from the exponential, persister, and VBNC cultures and washed by PBS twice and plated onto agarose gel pads for microscopy. For the VBNC culture (2 weeks or 5 weeks), cells were harvested by centrifugation and resuspended in LB containing ampicillin for 1 h or 3 h to remove antibiotic-sensitive cells before the PBS wash. The agarose gel pads were prepared with LB with 1.5% low melting temperature agarose (Nusieve GTG agarose – BMB # 50081); the melted LB agarose was poured into a template made from five glass slides (75 x 25 x 1 mm). The pad was covered by another slide glass and held together with a 50 g weight for 30 min to solidify. Samples (5 μL) were added to the agarose gel pad, covered by a coverslip, and sealed by nail polish to prevent evaporation. Cell growth at 37 °C on the agarose gel pad was observed every 20 min for up to 24 h by light microscopy (Zeiss Axioscope.A1, bl_ph channel at 1,000 ms exposure). During the observations, the microscope was placed in a vinyl glove box (Coy Labs) warmed by an anaerobic chamber heater (Coy Labs, 8535-025) to maintain 37 °C.

### Transmission electron microscopy (TEM) imaging

Samples (1 mL) of exponential and persister cells (3 h and 24 h) were pelleted by centrifugation at 17,000 x g for 2 min, washed twice with 1 mL of normal saline, and resuspended in 1 mL of normal saline. For VBNC cultures (5 weeks), 10 mL of cells were treated with ampicillin (100 μg/mL) for 3 h at 37°C to remove exponentially-growing cells, pelleted, washed once with 1 mL of normal saline, and resuspended in 1 mL of normal saline (10-fold concentration). In brief, for double staining^43^, cells were pelleted and fixed with 2% glutaraldehyde in 0.1M sodium cacodylate buffer for 1 to 12 h. Pellets were washed three times for 5 minutes with 0.1 M sodium cacodylate followed by secondary fixation with 1% osmium tetroxide for 1 h in the dark at room temperature. Next, samples were washed three times for 5 min with 0.1 M sodium cacodylate and for 5 minutes with water followed by en bloc staining with 2% uranyl acetate for 1-12 h. Samples were then dehydrated by using a series of ethanol washes (50%, 70%, 85%, and 95%, 3 x 100%), washed three times for 5 min with acetone, and embedded in Epon-Araldite (Ted Pella Inc, Redding, CA, USA). After the blocks had cured, they were sliced using an ultramicrotome, and 70 nm thin slices were collected on TEM grids (formvar/carbon coated grid) and post-stained with uranyl acetate and lead citrate. Sample grids were imaged by a Tecnai G2 Lab6 TEM at 200 kv.

### Flow cytometry

With exponential cells, rifampicin-induced persister cells (3 h), and VBNC cells (5 weeks old), metabolic activity was measured by flow cytometry. Cells were pelleted and washed twice with PBS. For the VBNC culture, to remove the exponentially growing cells, ampicillin (100 μg/mL) was utilized for 3 h before staining. For the dead cells (negative control), one mL of exponential culture was centrifuged, resuspended in 70% isopropanol, and incubated for 1 h at room temperature. The redox sensor and PI stains (BacLight™ RedoxSensor™ Green Vitality Kit, Thermo Fisher Scientific Inc., Waltham, MA, USA) were used with samples incubated at 37 °C for 10 min with light protection. The fluorescence signal was analyzed by flow cytometry (Beckman Coulter FC500) using the FL1 and FL3 channels.

### Tracing morphological changes during persister aging

At each time (3, 6, 9, 24 h), 1 to 3 mL of exponential and rifampicin-induced persister cell cultures were harvested by centrifugation at 17,000 x g for 1 min and resuspended in 1 mL of PBS. Morphological changes were observed by utilizing FM 4-64 fluorescent dye (10 μg/mL, Thermo Fisher Scientific Inc.), which stains cellular membranes, via fluorescence microscopy for improving the resolution to allow better observation of the cell size (Zeiss AxioA1, Blight field, 1,000 ms exposure and FM 4-64 filter 10,000 ms exposure). Cell roundness was calculated by ImageJ as 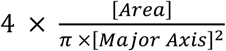.

## ACKNOWLEDGMENTS

This work was supported by the Army Research Office (W911NF-14-1-0279) and funds derived from the Biotechnology Endowed Professorship at the Pennsylvania State University. We appreciate the assistance with TEM provided by Missy Hazen (Microscopy and Cytometry Facility) and Jennifer Grey (Materials Research Institute) of the Penn State. We also thank Brian Dawson (Microscopy and Cytometry Facility, Penn State) for his assistance with flow cytometry.

## SUPPLEMENTARY VIDEO LEGENDS

### Supplementary Video 1. Cell division of exponentially-growing cells on an agarose gel pad

Exponential-phase cells growing on an agarose gel pad at 37 °C demonstrating that all cells grow immediately by cell division. Images were taken every 20 min, and the scale bar indicates 10 μm.

### Supplementary Video 2. Persister cell resuscitation on an agarose gel pad

Resuscitation of rifampicin-induced persister cells on an agarose gel pad at 37 °C demonstrating immediate resuscitation. Green arrows indicate cells that resuscitate immediately and blue arrows indicate cells that have delayed resuscitation. Images were taken every 20 min, and the scale bar indicates 10 μm.

### Supplementary Video 3. VBNC cell resuscitation on an agarose gel pad

Resuscitation of VBNC cells (5 weeks) on an agarose gel pad at 37 °C demonstrating no resuscitation. Images were taken every 1 hour, and the scale bar indicates 10 μm. Red arrows indicate spherical cells that did not resuscitate.

## SUPPLEMENTARY INFORMATION

**Supplementary Figure 1.**
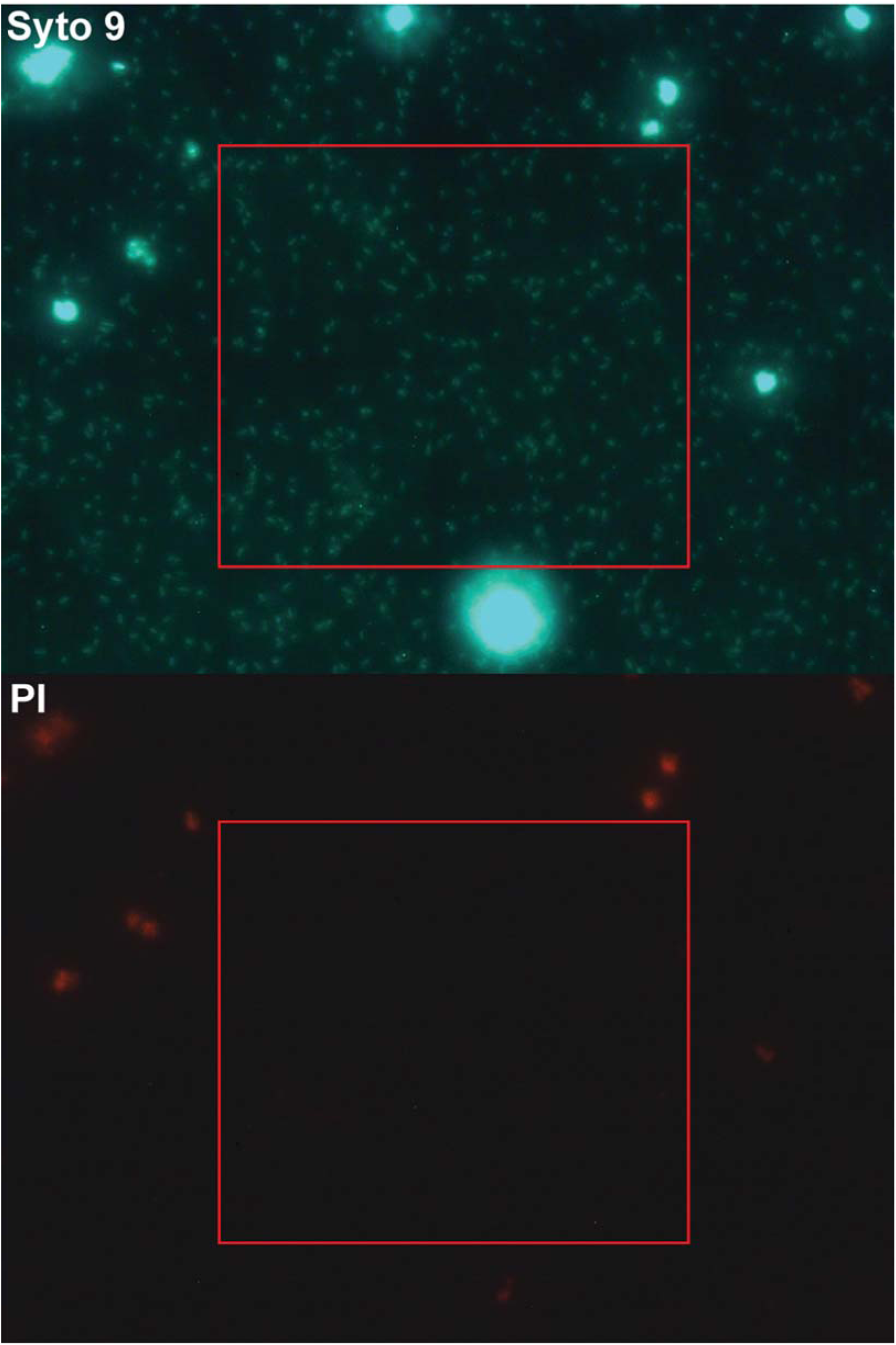
Counting total cell numbers in VBNC cultures using a hemotycometer. The cells in 19 squares (0.05 mm* 0.05 mm * 0.1 mm = 2.50E-04 μL) were counted using Syto 9 and PI staining. Total cell number was averaged based on the equation = [(Average cell number in the square * 1000) x 2.50E-04] * dilution factor.

**Supplementary Figure 2.**
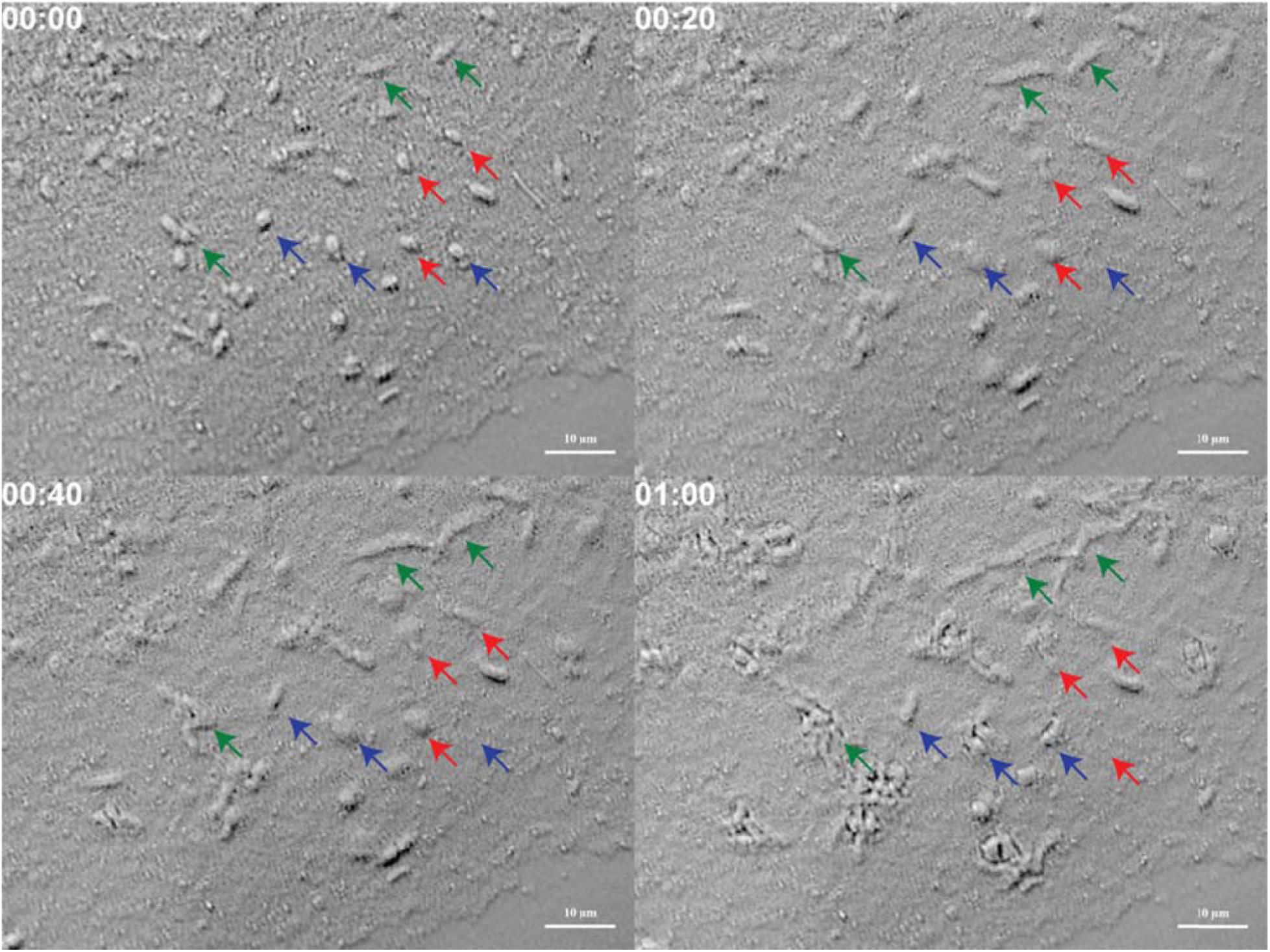
Resuscitation of morphologically-different VBNC cells (2 week old, ampicillin-treated at 100 μg/mL for 1 h). 2 week old cells formed under VBNC-inducing conditions were treated with ampicillin to lyse non-persister cells. The rod-shape cells (green arrows) resuscitated, some spherical cells (red arrows) did not resuscitate, and some spherical cells (blue arrows) resuscitated. Scale bar indicates 10 μm.

**Supplementary Figure 3.**
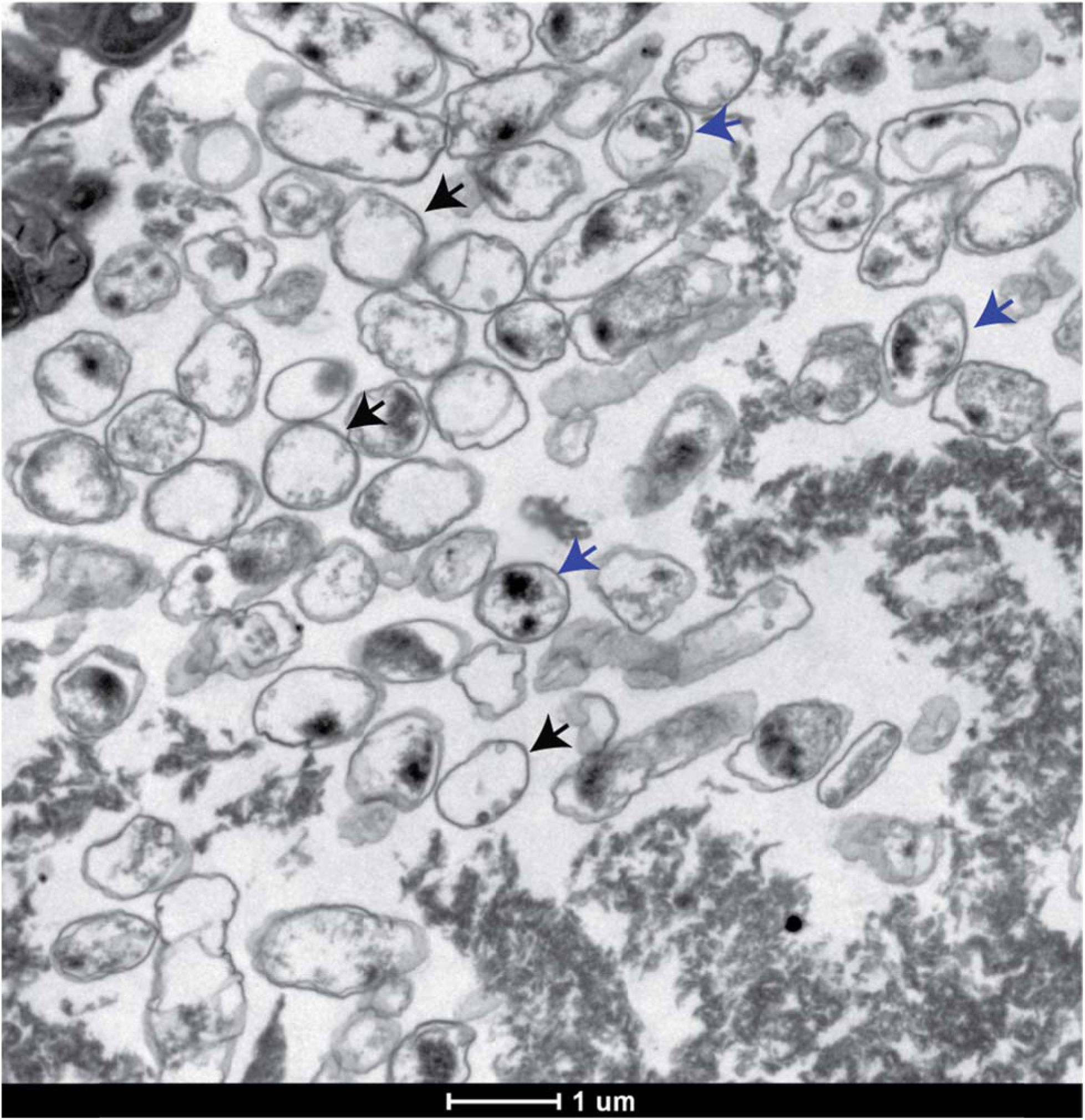
TEM image of a 5 week-old VBNC culture (ampicillin-treated for 3 h). Exponentially-growing cells were harvested and re-suspended in saline solution (0.85%) and cultured for 5 weeks at 37 °C, 250 rpm. Most VBNC cells had empty cytosol (black arrows) and some of the population had cytosol with cellular material reduced (blue arrows). Scale bar indicates 1 μm.

**Supplementary Figure 4.**
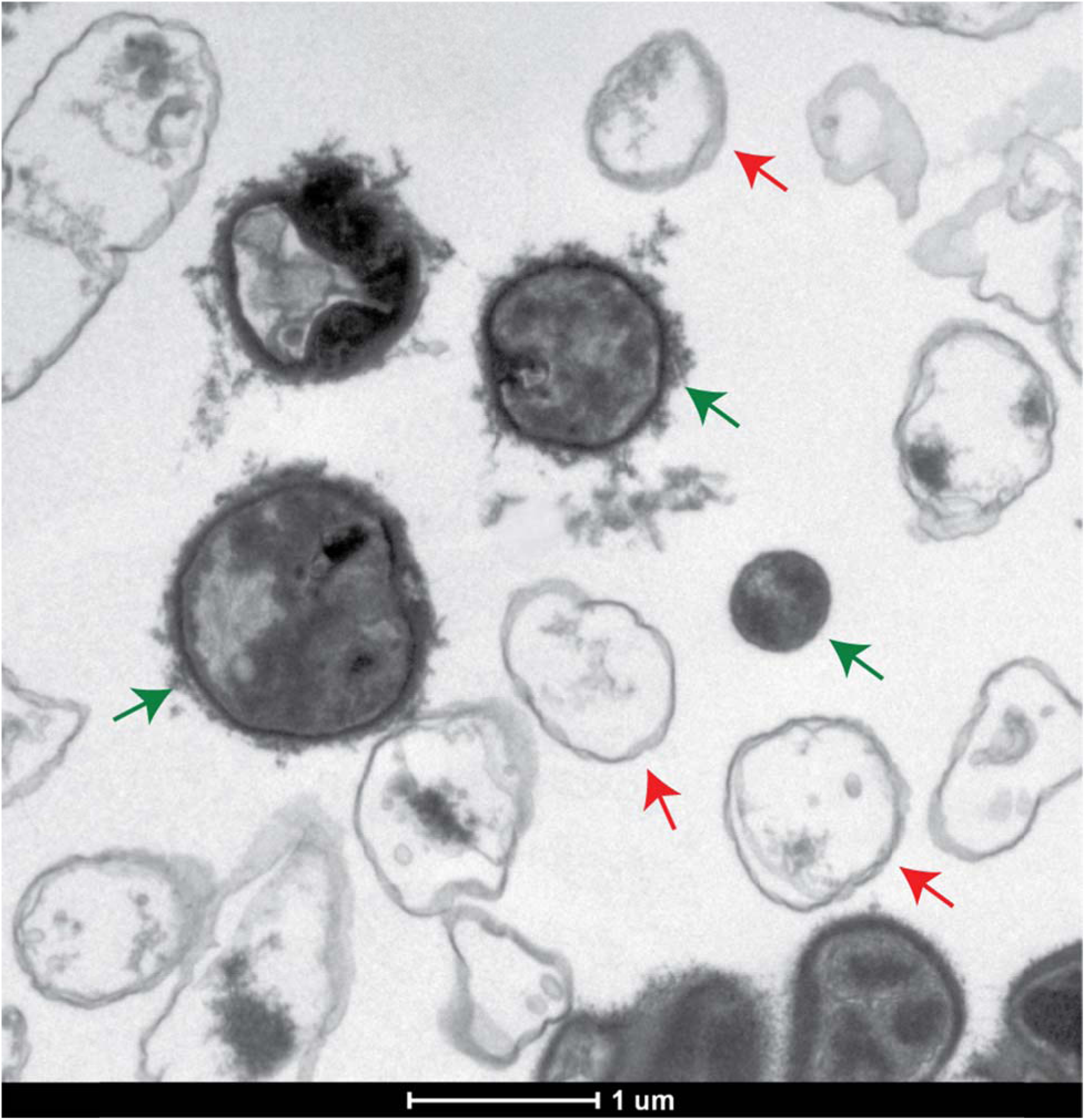
Two types of spherical cells in VBNC cultures (2 week old, ampicillin-treated at 100 μg/mL for 3 h). Green arrows indicate spherical cells with dense cytosol. Red arrows indicate spherical cells with empty cytosol. Scale bar indicates 1 μm.

**Supplementary Figure 5.**
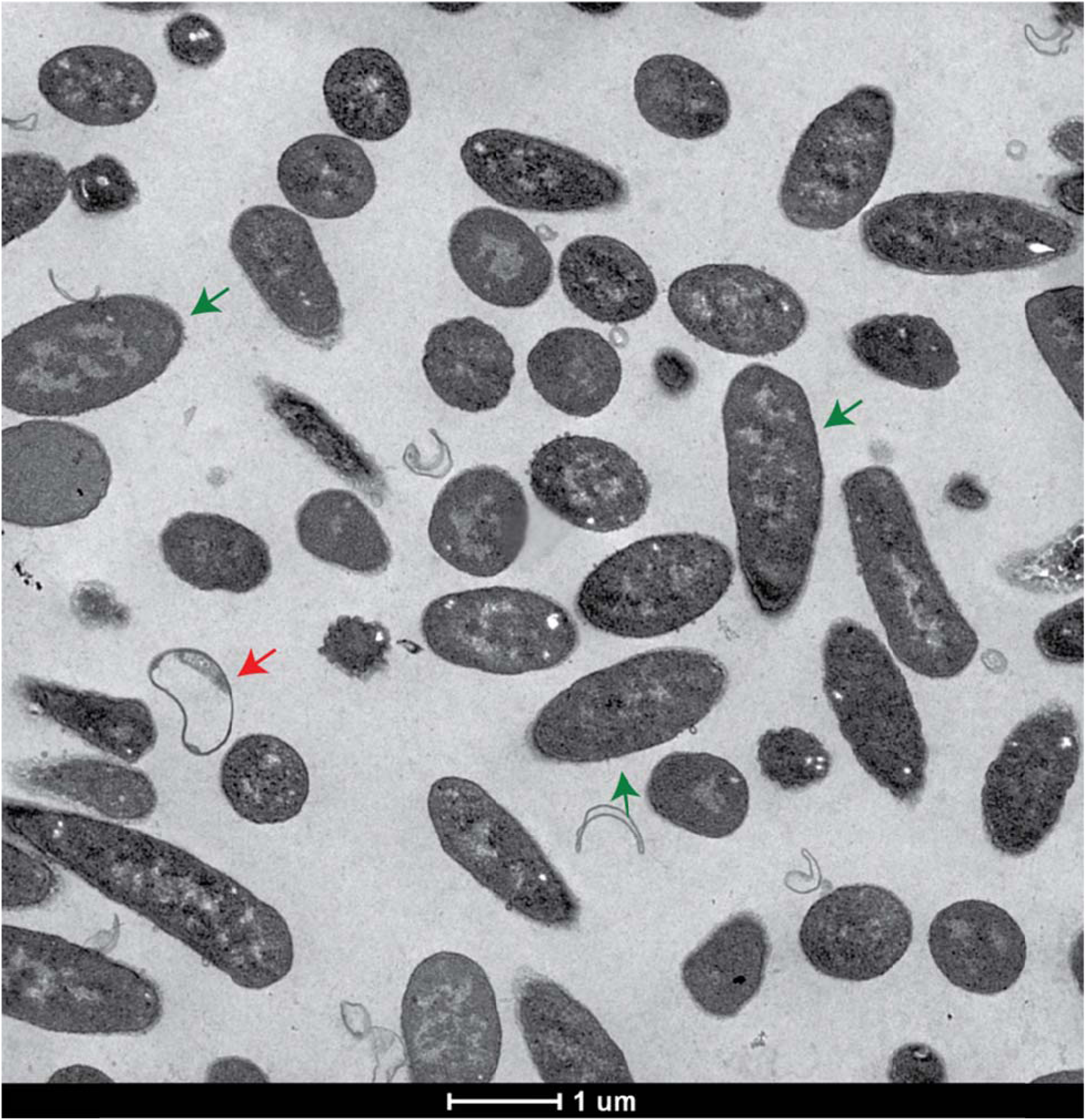
TEM image of fresh persister cells after 3 h in LB medium with ampicillin (100 μg/mL). Exponentially-growing cells were treated by rifampicin to form persister cells and ampicillin was used for 3 h to lyse non-persister cells. Most persister cells had dense cytosol and intact membranes (green arrows) while a few of the cells had VBNC-like empty cytosol (red arrow). Image is a larger field of view for the cells of Fig. 2. Scale bar indicates 1 μm.

**Supplementary Figure 6.**
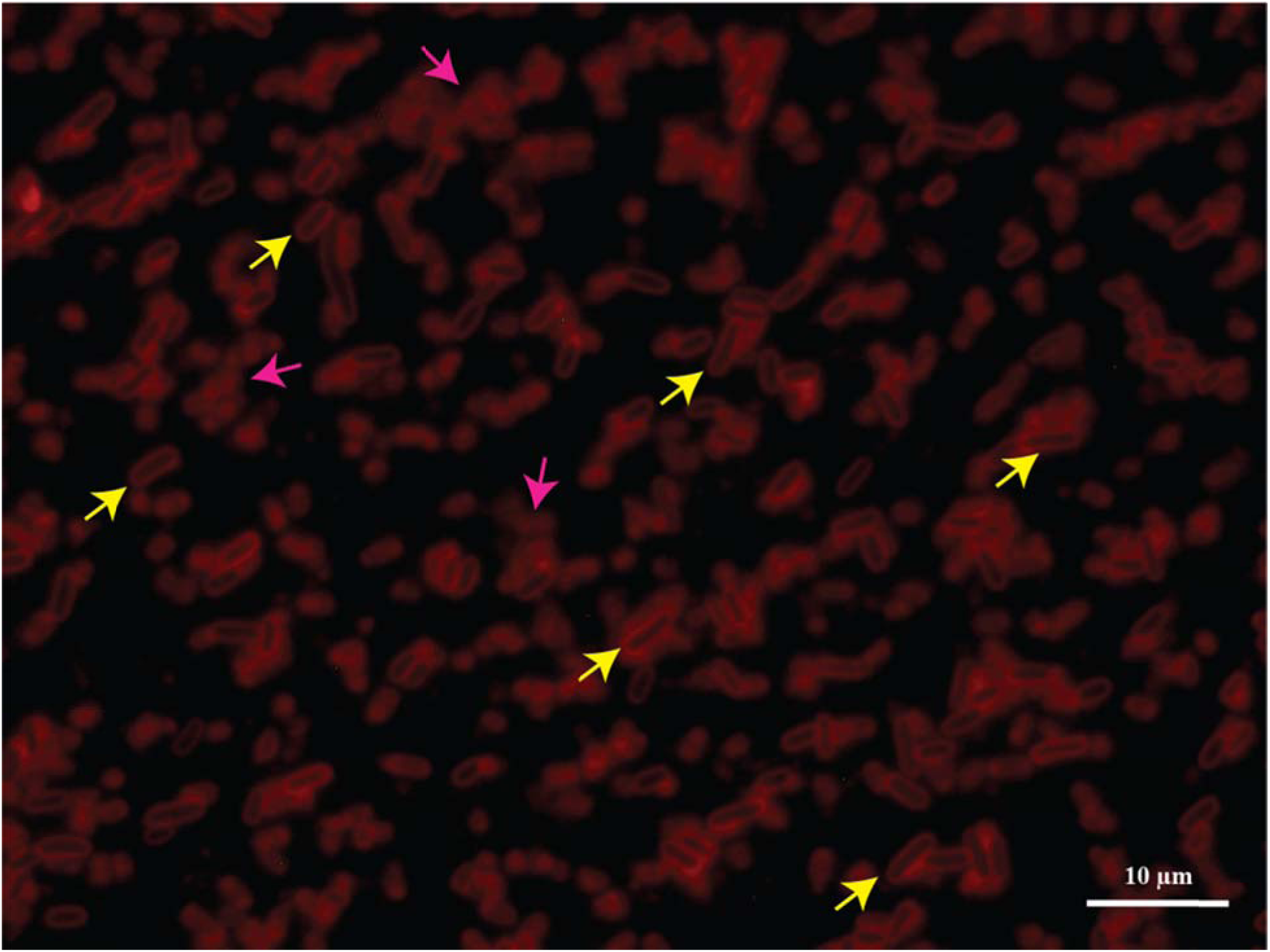
Persister cell morphology after 24 h in LB medium with ampicillin (100 μg/mL). Rifampicin-induced persister cells were cultured 24 hours in the presence of ampicillin and observed via fluorescence microscope by staining with membrane dye FM 4-64. Yellow arrows indicate representative cells that survive after the 24 h ampicillin treatment, and pink arrows indicate membrane-related debris. Scale bar indicates 10 μm.

**Supplementary Figure 7.**
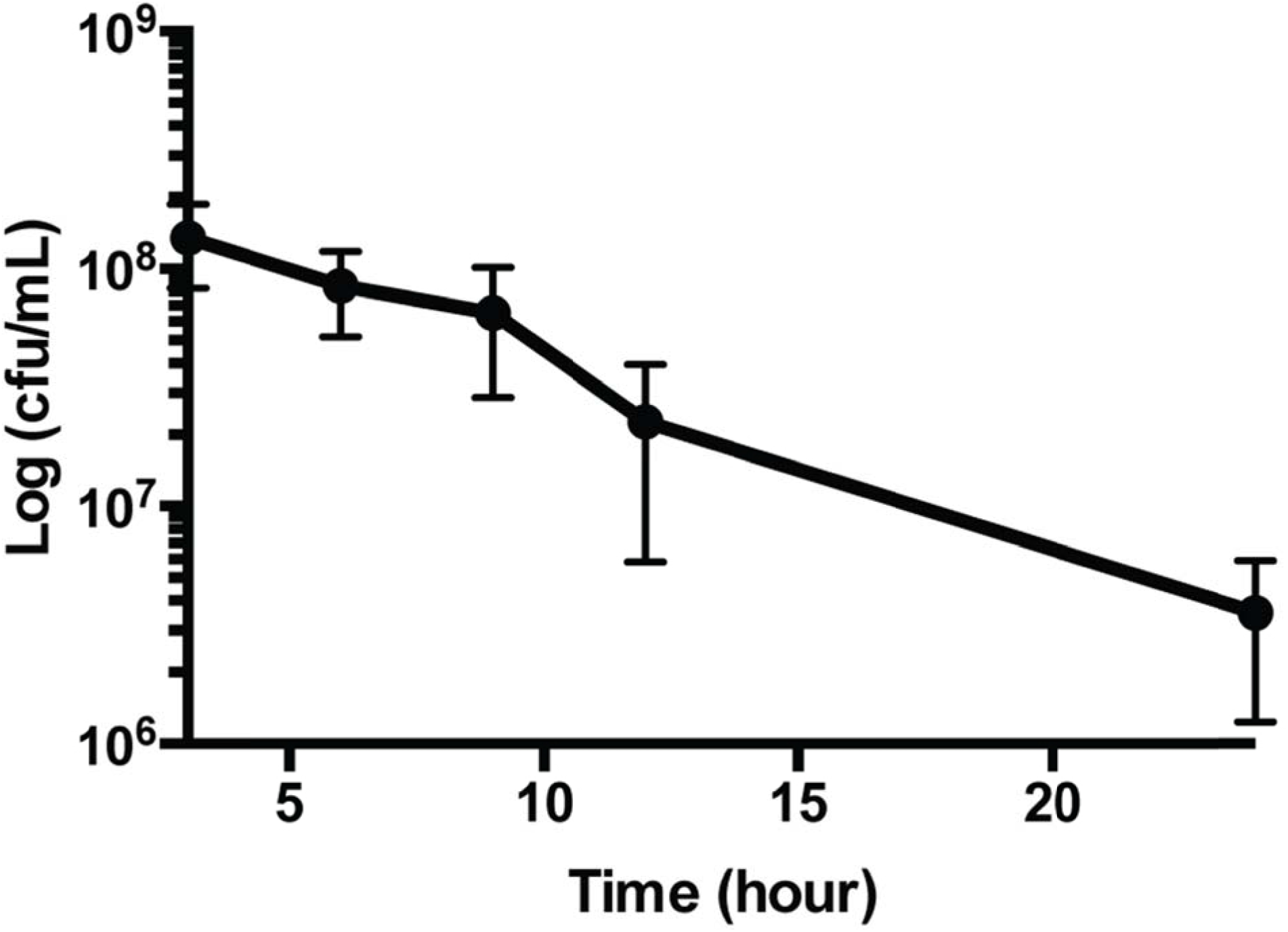
Persister cell viability during aging. The viability of rifampicin-induced persister cells in the presence of ampicillin was determined by plating. After 24 h, the viable persister cell number was reduced 70-fold, which is much lower than the cell death of exponentially-growing cells (reduction of 10^6^-fold in 24 h, data not shown).

**Supplementary Figure 8.**
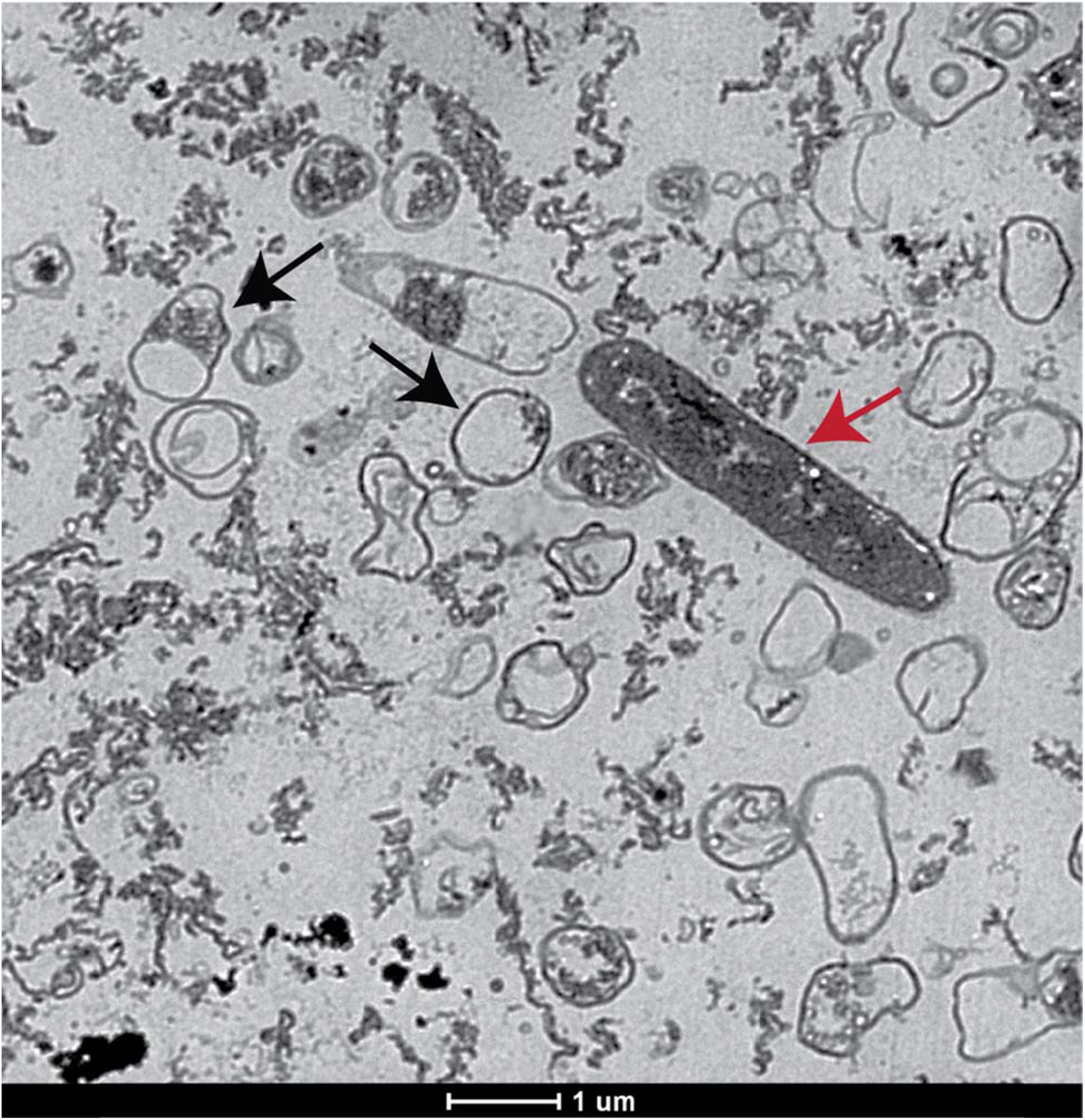
TEM image of aged persister cells after 24 h in LB medium with ampicillin (100 μg/mL). Exponentially-growing cells were treated by rifampicin to form persister cells and ampicillin was used to lyse non-persister cells. Most cells (black arrows) had empty cytosol and intact or damaged membranes like the VBNC culture. A few cells had dense cytosol and were rod-shape (red arrow). Scale bar indicates 1 μm.

**Supplementary Figure 9.**
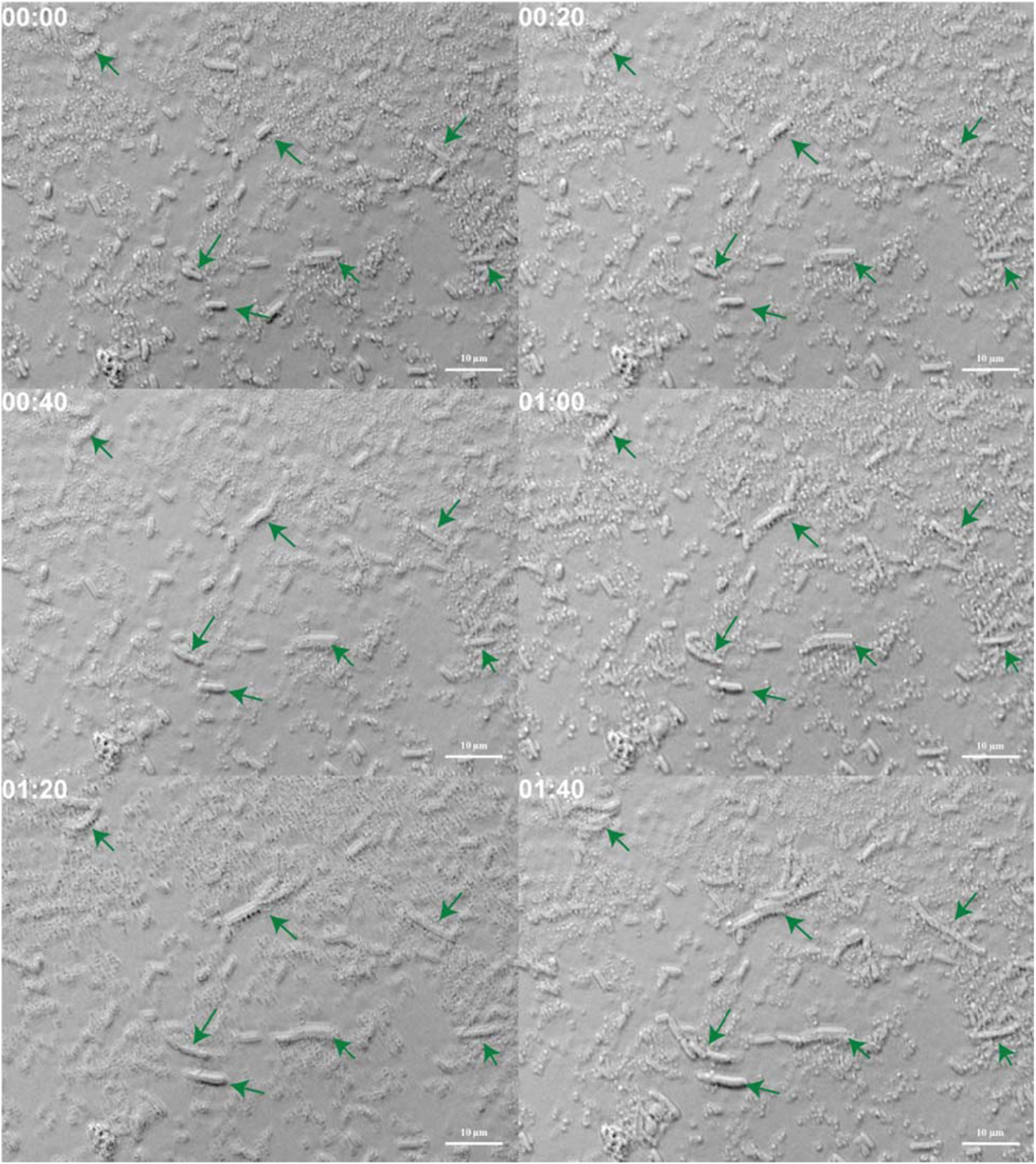
Resuscitation of aged persister cells after 24 h in LB medium with ampicillin (100 μg/mL). Exponentially-growing cells were treated by rifampicin to form persister cells and ampicillin was used to lyse non-persister cells. The surviving cells were rod-shaped (green arrows), and they resuscitated immediately by cell elongation then cell division. Scale bar indicates 10 μm.

